# Systematic analysis of CD39, CD103, CD137 and PD-1 as biomarkers for naturally occurring tumor antigen-specific TILs

**DOI:** 10.1101/2021.03.29.437255

**Authors:** Monika A. Eiva, Dalia K. Omran, Jessica Chacon, Daniel J. Powell

**Affiliations:** Ovarian Cancer Research Center, Department of Obstetrics and Gynecology, Perelman School of Medicine, University of Pennsylvania, Philadelphia, PA, United States; Center for Cellular Immunotherapies, Abramson Cancer Center, University of Pennsylvania, Philadelphia, PA, United States; Department of Pathology and Laboratory Medicine, Abramson Cancer Center, Perelman School of Medicine, University of Pennsylvania, Philadelphia, PA, United States; Texas Tech University Health Sciences Center, Paul L Foster School of Medicine and Woody L. Hunt School of Dental Medicine, El Paso, TX, United States

**Keywords:** CD137, tumor-infiltrating lymphocytes, tumor-specific biomarkers, CD103, PD-1, CD39

## Abstract

The detection of tumor-specific T cells in solid tumors is integral to the interrogation of endogenous antitumor responses and to the advancement of downstream therapeutic applications, such as checkpoint immunotherapy and adoptive cell transfer. A number of biomarkers are reported to identify endogenous tumor-specific tumor infiltrating lymphocytes (TILs), namely CD137, PD-1, CD103, and CD39, however a direct comparison of these molecules has yet to be performed. Here, we evaluate these biomarkers in primary human high-grade serous ovarian tumor samples using single-cell mass cytometry to characterize and compare their relative phenotypic profiles, as well as their response to autologous tumor cells *ex vivo*. CD137+, PD-1+, CD103+, and CD39+ TILs are all detectable in tumor samples with CD137+ TILs being the least abundant. PD-1+, CD103+, and CD39+ TILs all express a subset of CD137+ cells, while CD137+ TILs highly co-express the aforementioned markers. CD137+ TILs exhibit the highest expression of cytotoxic effector molecules, such as IFNγ and Granzyme B, compared to PD-1+, CD103+ or CD39+ TILs. Removal of CD137+ TILs from PD-1+, CD103+, or CD39+ TILs results in lower secretion of IFNγ in response to autologous tumor stimulation, while CD137+ TILs highly secrete IFNγ in an HLA-dependent manner. CD137+ TILs exhibited an exhausted phenotype with CD28 co-expression, suggestive of antigen recognition and receptiveness to reinvigoration via immune checkpoint blockade. Together, our findings demonstrate that the antitumor abilities of PD-1+, CD103+, and CD39+ TILs are mainly derived from a subset of TILs expressing CD137, implicating CD137 is a more selective biomarker for naturally occurring tumor-specific TILs.

## Introduction

The intratumoral abundance of tumor-infiltrating lymphocytes (TILs) is a positive prognostic factor for increased survival in most solid cancers, indicating that TILs are integral to endogenous antitumor immunity and play a role in controlling cancer progression^1,2^. However, only a small percentage of TILs respond against tumor antigens and their antitumor response can be hindered by multiple mechanisms of immunosuppression^3,4^. The challenges of detecting TILs capable of responding to tumor antigens has led to great interest in identifying biomarkers of tumor-specific TILs in solid tumors. Biomarkers that identify tumor-specific TILs are integral for downstream applications, such as enriching tumor-specific TILs for use in adoptive cellular therapy, investigating endogenous antitumor immunity, studying mechanisms of effective immunotherapy, identifying antigen-specific T-cell receptors or neoantigens, and exploring the immunobiology of these cells^5–8^. The need for effective biomarkers to detect T cells is further underscored by the fact that many cancers, such as ovarian cancer, do not have well-defined shared tumor-specific antigens capable of initiating a tumor-specific T cell response. The lack of shared tumor-specific antigens is in contrast to other cancers, such as melanoma, where some patients mount spontaneous responses against the melanocyte differentiation antigen, MART-1, which can be used to rapidly identify tumor-specific T cells in melanoma patients using peptide/MHC detection agents^9^. Furthermore, many cancers including ovarian cancer, have limited numbers of T cells that naturally respond to tumor-specific antigens, making their examination challenging. Identifying robust biomarkers for tumor-specific TILs can address this issue.

Various biomarkers are used to detect endogenous tumor-specific T cells from solid tumors, such as the co-stimulatory receptor CD137 (also known as 4-1BB and TNFRSF9), the negative immunoregulatory receptor PD-1, the lymphocyte-retention mediating integrin CD103, and the co-expression of both the ectonucleotidase CD39 and CD103^10–13^. Identifying a singular, accurate biomarker for tumor-specific TILs would streamline downstream research and clinical applications, but it is unknown which singular biomarker is most effective at identifying the tumor-specific TIL subset, as a direct comparison of these reported biomarkers has not been performed. Addressing this knowledge gap is particularly important, because TILs frequently co-express these markers and each biomarker can be differentially expressed across the TIL population, therefore, a biomarker comparison is needed to identify the marker that most accurately discerns tumor-specific TILs^14,15^.

Here, we compared the expression of CD137, PD-1, CD103 and CD39 on TILs in human ovarian cancer, as these are leading biomarkers used to identify tumor-specific TILs. We hypothesized that a comparative interrogation of TILs in human tumors would reveal which biomarker is most discriminating for tumor-specific TILs with autologous antitumor activity.

## Materials and methods

### Tumor Samples

Viably frozen, human high-grade serous ovarian tumor samples were purchased from the Penn Ovarian Cancer Research Center (OCRC) Tumor BioTrust Collection. Ethics statement: All donor samples used in this study were de-identified and approved for use by the UPenn Institutional Review Board (IRB 702679, UPCC 17909). Sex and weight are not a biological variable as all tumor samples are from females. As samples are de-identified, age and weight are not known. Surgically resected tumors were procured from the operating room in an aseptic manner. Tissue was mechanically processed into fragments and added to an enzyme digest solution. A 10X stock solution of the enzyme digest buffer contains 2 mg/ mL collagenase (Sigma Aldrich) and 0.3kU/mL DNase I Type IV (Sigma Aldrich); solution was diluted to a 1x solution with RPMI 1640 at time of digestion. Tissue was incubated in the enzyme digest buffer overnight at room temperature on a rotator. Dissociated tumor tissue was subsequently filtered through sterile 100μm nylon mesh, centrifuged, and washed twice with dPBS (Dulbecco’s Phosphate Buffered Saline). Resultant tumor cell digests were cryopreserved in 10% dimethyl sulfoxide (DMSO) (Sigma Aldrich) and human serum (Valley Biomedical, Inc., Product #HS1017). Samples were frozen at −80C and banked at - 150C until further use.

### Mass Cytometry staining

CyTOF antibodies were bought from Fluidigm as pre-conjugated metal tagged antibodies or were conjugated in-house using the Maxpar Fluidigm kit and protocol. All antibodies were titrated to determine optimal concentrations for staining samples. The panel used to initially investigate tumor-specific markers, before inclusion of CD39 in the aforementioned panel, had the following surface markers: CD3, CD45, CD4, CD8, CD244, CD69, OX40, Lag-3, CD103, Tim-3, TIGIT, PD-1, CD137, CD28, CD127, CD27, GITR, CD25, HLA-DR, and CD160. Intracellular antibodies included: CTLA-4, pStat5, IL-17A, IL-2, IFNg, Granzyme B, Ki67, and Perforin. We subsequently designed a panel to include CD39 and all other tumor-specific markers of interest. The following panel included CD39 and was used for downstream viSNE, metaPhenoGraph, and biaxial analysis. Anti-human surface markers for the panel were: CD3, CD45, CD4, CD8, CD103, PD-1, OX40, CD39, CD69, CD25, CD137, CD27, Tim-3, CD127, CD28, CD244, CD5, Lag-3, TIGIT, HLA-DR, and CD160. Intracellular markers included: Ki67, IL-17A, IL-2, IFNg, IL-6, Perforin, pStat5, TNFa, Granzyme B, CTLA-4, and EOMES. The initial panel to compare CD39 and CD137 positive TILs had the following surface antibodies interrogated: CD3, CD45, CD4, CD8, CD137, CD39, CD25, HLA-DR, and CD127. Intracellular antibodies detected for were: IL-2, pStat5, EOMES, T-bet, IL-17A, IFNg, Granzyme B, Ki67, and Perforin. The last panel used in this study was designed to focus on TIL exhaustion. Surface antibodies used were: CD3, CD45, CD4, CD8, OX40, CD103, TIGIT, CD137, CD39, CD25, CD3, HLA-DR, and CD127. Intracellular antibodies were: IL-2, pStat5, EOMES, T-bet, and Ki67. For all panels, cell identifier stain Iridium191/193, live identifier 127IdU (Fluidigm) were used. To discriminate dead cells, cisplatin purchased from Fluidigm or dead stain maleimido-mono-amine-DOTA (mm-DOTA) from Macrocyclics was used. Viably frozen ovarian human tumor digests were stained for CyTOF following the same methodology as Bengsch et al., 2018 ^16^. Data acquisition was performed on a CyTOF Helios (Fluidigm CyTOF Helios Mass Cytometer, RRID:SCR_019916) by the CyTOF Mass Cytometer Core at UPenn. The core performed bead-based normalization for all samples.

### Fluorescent-Activated Cell Sorting

Tumor samples were thawed and washed twice with staining buffer (phosphate-buffered saline, 5% fetal bovine serum) to remove DMSO. Samples were subsequently stained with Zombie aqua (BioLegend Cat# 423102) for 10 minutes to discriminate live and dead cells. Samples were washed twice to remove Zombie aqua, then incubated at 4°C for 30 minutes in 50ul of an antibody cocktail to label human surface markers. Following surface staining, samples were washed three times. Samples were sent to the Flow Cytometry Facility at the Wistar Institute for fluorescent-activated cell sorting (FACS) on a MoFlo Astrios or to the Flow Cytometry Core at the Children’s Hospital of Philadelphia and sorted on an Aria. All antibodies were purchased from BioLegend. For all analyses, singlets were detected using FSC-H versus FSC-A followed by SSC-H versus SSC-A. Cells negative for Zombie aqua, were identified as live cells. Anti-human-anti-CD3-PerCpCy5.5 (BioLegend Cat# 317336, RRID:AB_2561628) was used to detect T cells and the following anti-human antibodies were used to identify T cell subsets: anti-CD137-PeCY7 (BioLegend Cat# 309818, RRID:AB_2207741), anti-CD103-BV605 (BioLegend Cat# 350218, RRID:AB_2564283), anti-CD39-APC (BioLegend Cat# 328210, RRID:AB_1953234), and anti-PD-1-APCCy7 (BioLegend Cat# 329922, RRID:AB_10933429).

### Mass cytometry biaxial analyses

Traditional biaxial analysis, on bead-normalized fcs files, was performed using Flowjo V10 software (FlowJo, RRID:SCR_008520). Intact single cells were identified using event-length and Iridium. Cells were live-gated according to 127IdU and mm-DOTA, where dead cells are positive for mm-DOTA. CD3 and CD45 positivity identified T-cells. Sequential gating analysis was performed for all analyzed markers. The resulting values were used to determine population frequencies.

### viSNE, and metaPhenoGraph analyses

High-dimensional analysis was conducted using the algorithm viSNE, which uses the Barnes-Hut t-SNE (bh-SNE) implementation, from *cyt* a visualization tool written in Matlab (R2016b, MATLAB,RRID:SCR_001622) downloaded in 2015 and available at https://www.c2b2.columbia.edu/danapeerlab/html/cyt-download.html. Live, single, CD3^+^CD45^+^CD137^+/-^ exported fcs data from five donor samples were imported into *cyt*, arcsinh5-transformed, and run as described by Amir et al., 2013 ^17^ to create viSNE plots. The following parameters were used for bh-SNE mapping analysis: Ki67, IL-17A, IL-2, IFNγ, CD103, PD-1, IL-6, OX40, CD39, Perforin, CD69, CD4, CD8, pStat5, TNFa, GITR, CD25, Granzyme B, CD137. The PhenoGraph algorithm was run, as described by Levin et al., 2015 ^18^, with a nearest neighbor input of k=30 and a Euclidean distance metric. Markers used for PhenoGraph clustering were the following: Ki67, IL-17A, IL-2, IFNg, CD103, PD-1, IL-6, OX40, CD39, Perforin, CD69, CD4, CD8, pStat5, TNFa, GITR, CD25, Granzyme B, CD137. PhenoGraph was metaclustered, as described by Levine et al., 2015, using a k=15 and a Euclidean distance metric. viSNE, PhenoGraph, metaPhenoGraph plots and heatmaps were created by *cyt*.

### Co-culture experiment

The following T cell subsets were FACS sorted from patient tumor samples: CD137^+^, CD39^+^CD137^-^, CD103^+^CD137^-^, and PD-1^+^CD137-using the BioLegend antibodies specified in the FACs sorting section. T cells were rested overnight in media. CD45^+^ cells were depleted from the same patient sample to obtain CD45^-^cells for co-culture using the EasySep Human CD45 Depletion Kit from StemCell Technologies Cat# 17898. T cell subsets were co-cultured, with 10ug/ml HLA-blocking Class I (BioLegend Cat# 311402, RRID:AB_314871) & II (BioLegend Cat# 361702, RRID:AB_2563139) or isotype (BioLegend Cat# 400202) antibodies, at a 1:2 ratio of T cells to autologous tumor cells in 100ul of media in a 96-ubottom plate. Following 24hrs co-culture, samples were spun down at 1300rpm, and supernatants were collected and frozen at −80C. To analyze cytokines within the supernatants, the manufacture’s protocol of the LEGENDplex Human CD8/NK Panel kit (BioLegend Cat# 740267) was followed, and two technical replicates were analyzed per sample.

### Statistical analysis

The Student’s two-tailed, paired t-test was run to determine statistical significance. NS represents a p-value >0.050, (*) represents a p-value ≤0.050, (**) represents a p-value <0.01, (***) represents a p-value < 0.001, (****) represents a p-value < 0.0001, error bars represent 95% Confidence Interval.

## Results

### A subset of TILs express effector molecules

To investigate the phenotype of TILs harbored within infiltrated tumors, the algorithms viSNE and PhenoGraph metaclustering ^17,18^ were used to co-map CD3^+^CD45^+^ TILs in enzyme-digested ovarian tumors analyzed by single-cell mass cytometry. To address patient-specific variability and to understand TIL dynamics shared between samples, PhenoGraph clusters were merged using the metaclustering algorithm in the interactive *cyt* tool ^17^. Metaclustering analysis identified seven major TIL populations (**Figure 1A**). Metaclusters (MCs) 1, 5, 6, and 7 were generally conserved among all samples tested, while MCs 2, 3, and 4 had greater variability (**Figure 1B**). MC5 (mean= 1.51, 95% CI= −0.37 to 3.39) and MC6 (mean= 2.21, 95% CI= −0.83 to 5.24) were the rarest subsets in all samples, and MC5 was consistently enriched for cells expressing activation, proliferation and effector molecules (**Figure 1B**,**C**). Compared to the other metaclustered groups, only MC5 highly expressed effector molecules associated with antitumor responses, including IL-2, IFNγ, perforin, TNFα, and Granzyme B (**Figure 1C**).

**Figure 1.**
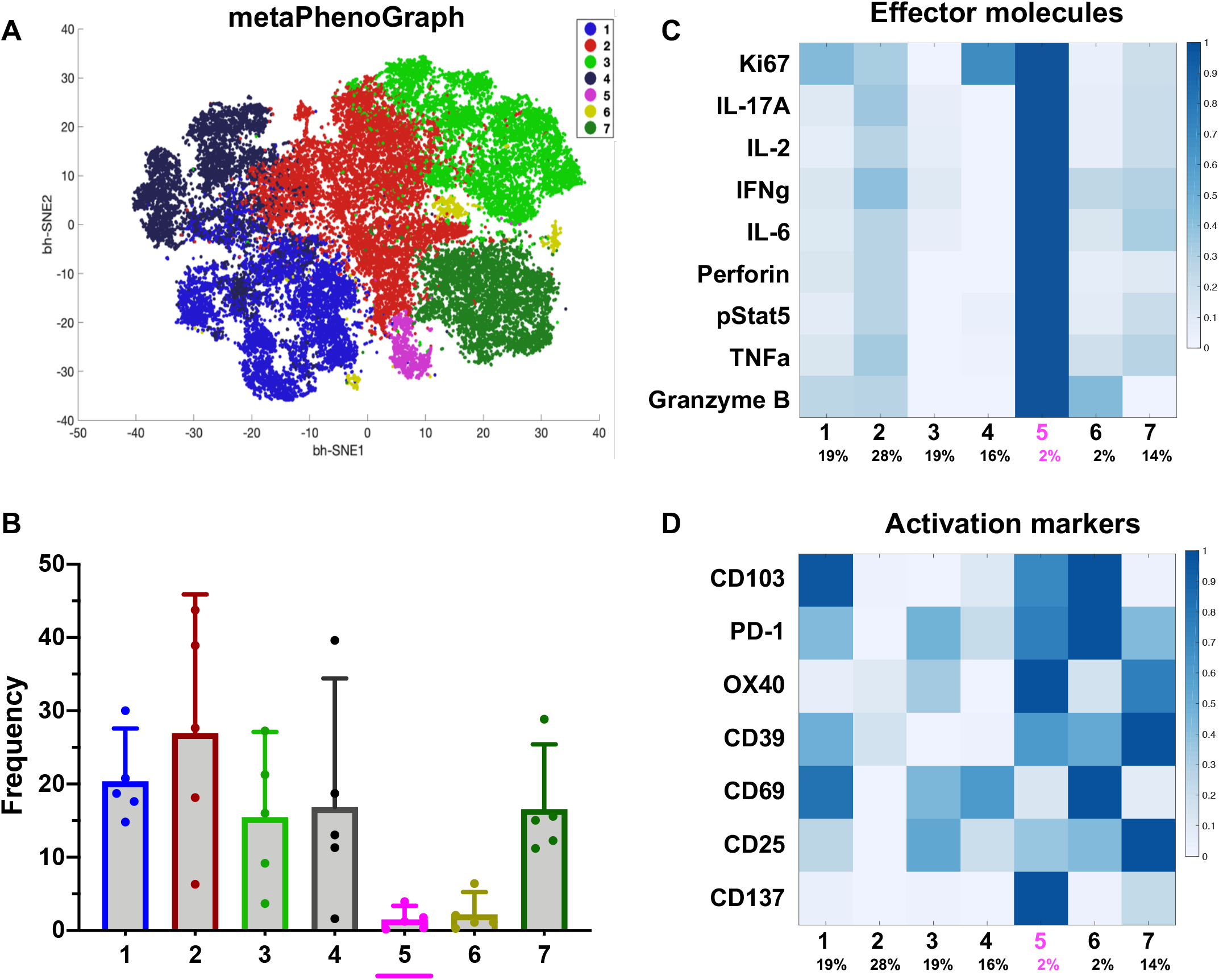

A series of activation-associated, cell surface markers have recently been described to identify, characterize and utilize naturally-occurring tumor-specific T cells in human tumors. CD137, PD-1, CD103, and CD39 are most commonly utilized as biomarkers of TILs with tumor-specificity^10–13,19^. MC5, which highly expresses effector molecules, was enriched for TILs expressing high levels of CD137 as well as the co-stimulatory receptor OX40, another TNFR family member upregulated upon T cell activation. MC5 moderately expressed CD103, PD-1, and CD39, as well as activation markers CD69 and CD25 (**Figure 1D**), indicating that cells in MC5 are enriched for an activated T cell population.

### CD137^+^ TILs preferentially express effector molecules and co-express biomarkers of tumor-specificity

To gain a further understanding of which biomarkers are most selective in identifying TILs expressing effector molecules within human cancer, we examined viSNE plots of activation and tumor-specific biomarkers, which revealed the heterogeneity of their expression patterns. Similar to what was observed in the metaPhenoGraph heat map results (**Figure 1D**), CD137 expression was primarily detected in the MC5 region, was expressed by both CD4^+^ and CD8^+^ TILs, and had co-expression of OX40, CD103, CD39, and PD-1 (**Figure 2A**). PD-1 and CD69 expression was common, broadly distributed, with overlapping expression of CD25, OX40, CD103, CD39, and CD137. CD25 and OX40 expression was dominated by CD4+ TILs and commonly co-expressed with CD39, while CD103+ TILs were mainly CD8+, a portion of which expressed CD39. Overall, few TILs expressed effector molecules, such as IFNγ, IL-2, and TNFα, and their MMI was low, compared to the level of activation and tumor-specific biomarkers. However, the few TILs that expressed effector molecules such as IFNγ, IL-2, and TNFα, were positive for CD137 in the MC5 region, suggestive of CD137^+^ TIL polyfunctionality, and CD137 expression was most focal than other tumor-specific biomarkers (**Figure 2A**).

**Figure 2.**
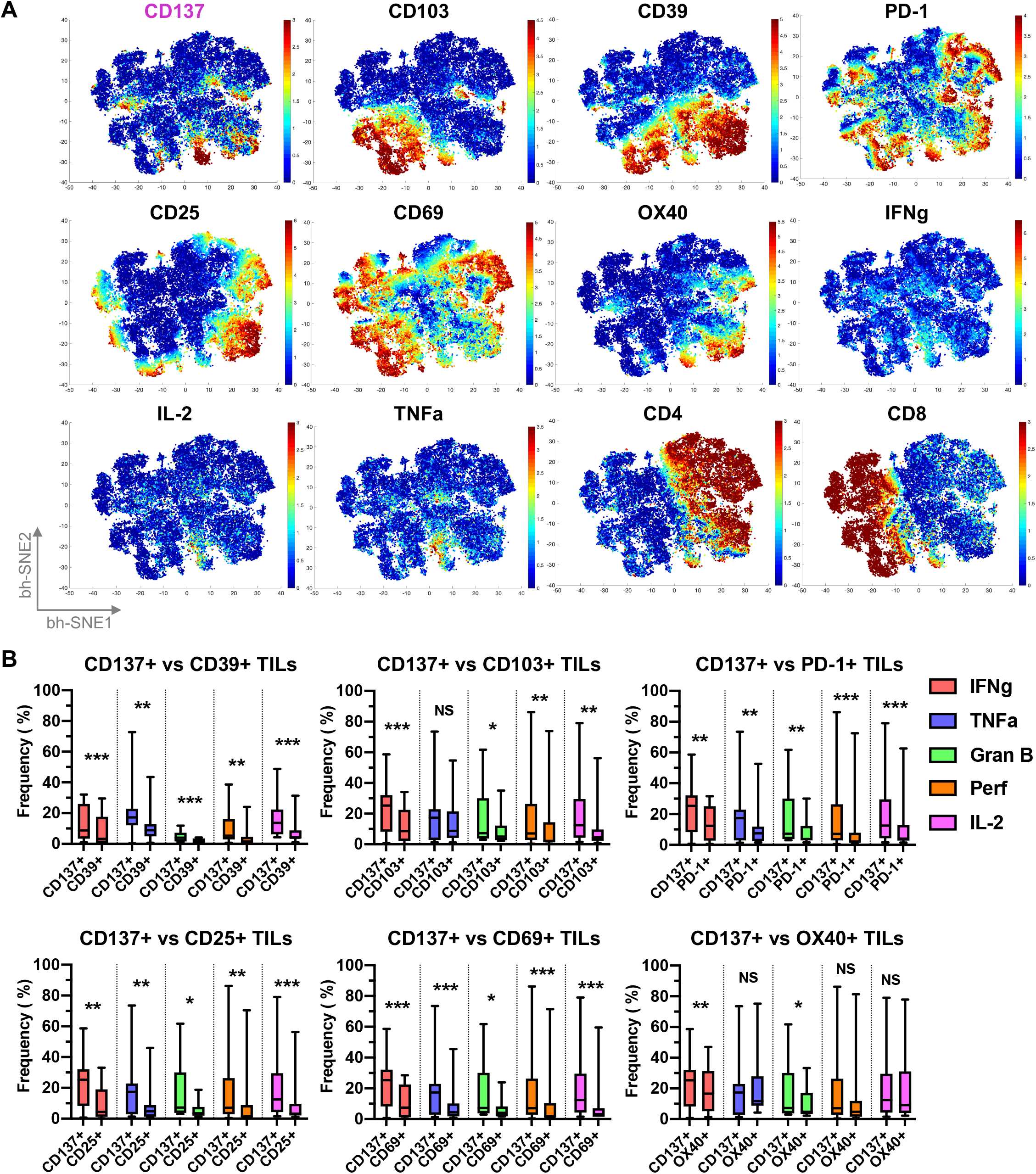

Since viSNE plot analyses indicated that CD137 expression overlapped more with effector molecule expression than other biomarkers, we compared effector molecule expression within the CD137^+^ TIL population to expression in TILs expressing other tumor-specific and activation markers (**Supplementary Figure 1**). Generally, CD137^+^ TILs exhibited the greatest frequency of cells expressing IFNγ, TNFα, Granzyme B, perforin and IL-2, compared to other biomarker expressing TILs (**Figure 2B**). CD137^+^ TILs had greater expression of IFNγ (p-value = 0.0008), Granzyme B (p-value =0.01), perforin (p-value = 0.003), and IL-2 (p-value = 0.002) than CD103^+^ TILs, but similar levels of TNFα expression (**Figure 2B**). CD137^+^ TILs and OX40^+^ TILs were similar with the exception of CD137+ TILs expressing greater frequencies of IFNγ (p-value = 0.008) and Granzyme B (p-value = 0.01) (**Figure 2B**). While this study focuses on comparing single biomarkers, dual expression of CD103^+^CD39^+^ was reported to identify CD8^+^ tumor-specific TILs ^13^. When comparing CD103^+^CD39^+^ TILs to CD137^+^ TILs, CD137 expression was more selective for identifying total CD3^+^ and CD8^+^ TILs expressing effector molecules (**Supplementary Figure 2 A**,**B**) with no differences observed when comparing CD4^+^ TILs (data not shown).

Although CD137^+^ TILs exhibited the highest expression of effector molecules, the frequency of these cells was low (mean =4.1%, 95% CI =1.87 to 6.36) compared to TILs expressing other biomarkers (**Figure 3A**). Since viSNE and PhenoGraph analyses (**Figure 1**) revealed that CD137^+^ TILs often co-express tumor-specific biomarkers, we next examined the frequency of CD137^+^ TILs within TIL populations expressing other tumor-specific biomarkers using biaxial gating (**Figure 3B**). CD137^+^ TILs commonly co-expressed PD-1 (54.9% 95% CI=39.47 to 70.31]), CD103 (mean = 37.6%, 95% CI=24.64 to 48.78), and CD39 (mean = 76.8%, 95% CI=64.55 to 88.95). In contrast, only a small portion of PD-1^+^ (mean = 6.2%, 95% CI=4.36 to 8.07), CD103^+^ (mean = 6.2%, 95% CI= 3.13 to 9.17), or CD39^+^ (mean = 6.7%, 95% CI=3.45 to 10.02) TILs co-expressed CD137. These results, combined with effector molecule expression data (**Figure 2**), indicate that CD137 is the more selective marker for identifying tumor-specific TILs (**Figure 3C**).

**Figure 3.**
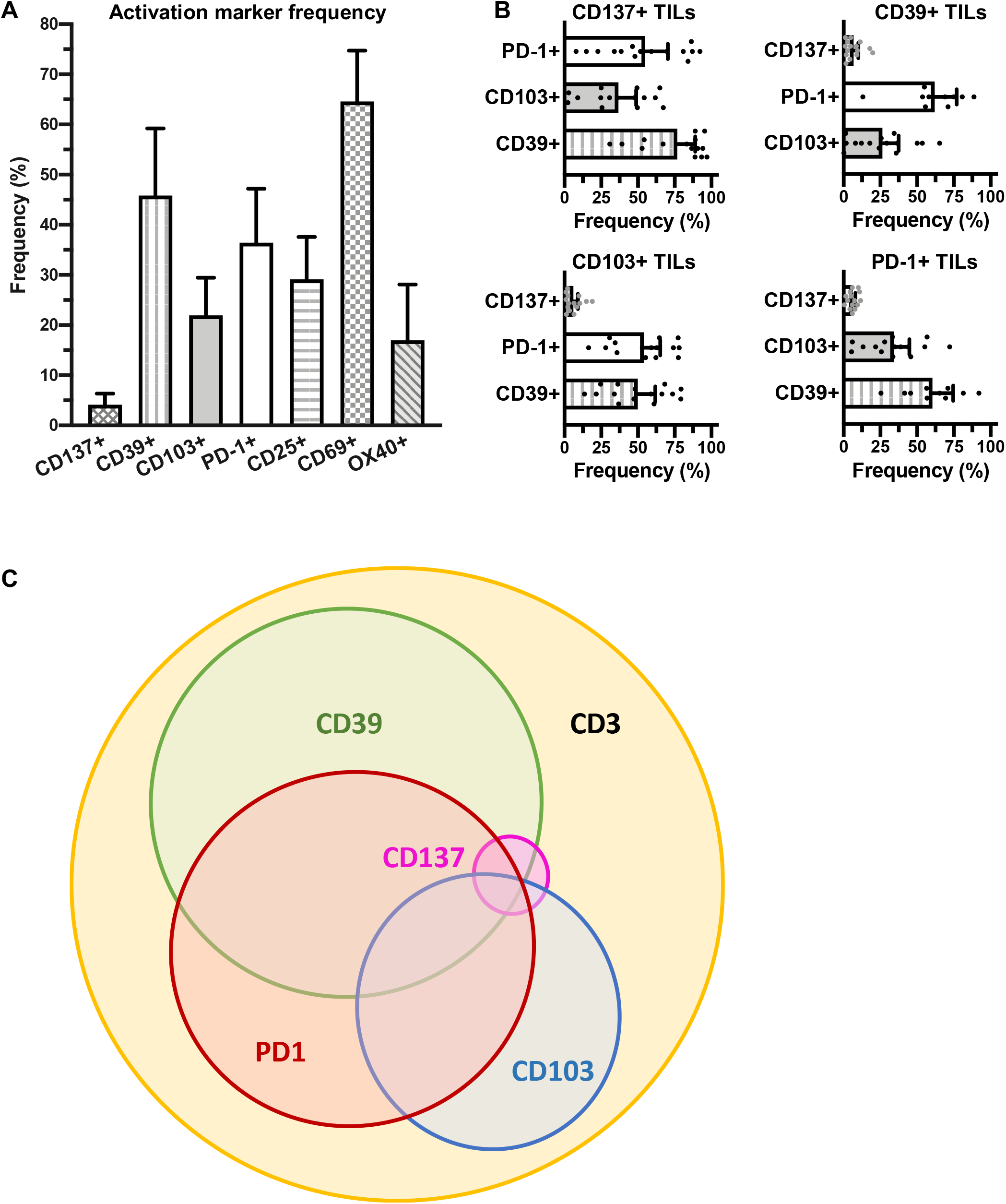

### CD137^+^ TILs are the subset of PD-1^+^, CD103^+^, and CD39^+^ TILs that express effector molecules and exhibit antitumor activity

We next investigated whether the CD137^+^ TIL subset contained within other biomarker populations are enriched for effector molecules. Decreased IFNγ expression was observed in CD39^+^ (*p*-value = 0.001), CD103^+^ (*p*-value< 0.001), and PD-1^+^ (*p*-value= 0.002) TILs when CD137^+^ TILs were selectively gated out (**Supplementary Figure 2C)** prior to analysis in **Figure 4A**. This effect was also observed in TILs expressing CD25 (*p*-value= 0.001), CD69 (*p*-value= 0.001), or OX40 (*p*-value= 0.003) (**Figure 4A**). Granzyme B expression similarly decreased (**Figure 4B**), leading us to hypothesize that the CD137^+^ TIL subset may account for the reactivity observed in other biomarker-expressing tumor-specific TIL populations ^10–13^.

**Figure 4.**
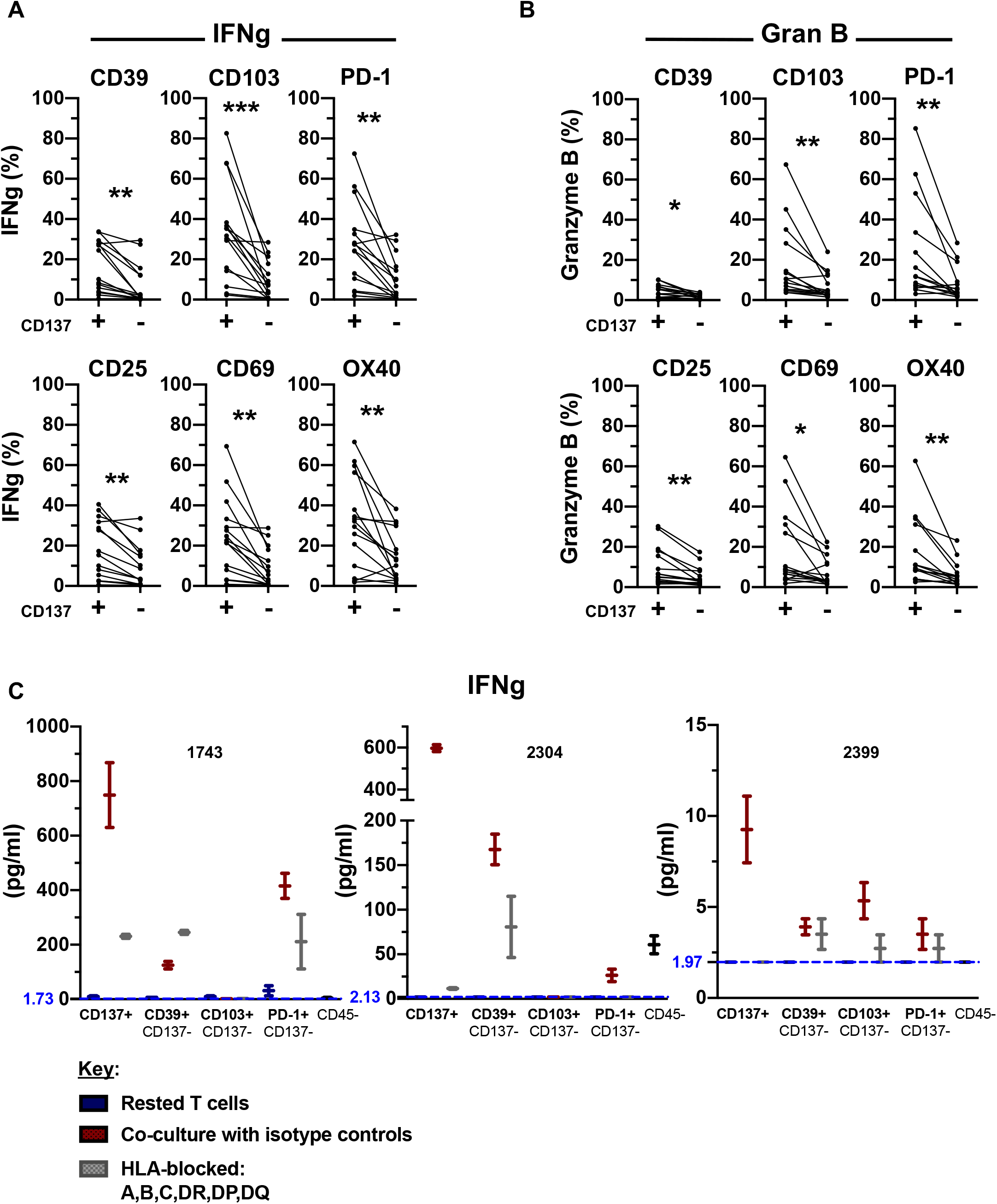

We next tested whether functional reactivity of TILs was restricted to the CD137^+^ TIL subset in co-culture assays where CD137^+^ TILs were first sorted out of the bulk TIL, and then other biomarker expressing TIL subsets were sorted prior to co-culture with autologous tumor cells (**Supplementary Figure 2D**). Compared to the PD-1^+^, CD103^+^ and CD39+ TIL populations depleted of CD137^+^ cells, the CD137^+^ TIL subset produced the highest levels of IFNγ in response to autologous tumor cells exposure in three independent donor samples. Production of IFNγ by CD137^+^ TILs upon autologous tumor co-culture was HLA-dependent, as IFNγ decreased upon HLA blocking of MHC class I and class II with antibodies (**Figure 4C**). In all tested TIL samples, the CD137^+^ subset secreted IFNγ levels twice as high as that of unstimulated TILs alone. These results indicate that effector molecule expression is enriched within the CD137^+^ TIL fraction, and that CD137^+^ TILs account for the majority of antitumor reactivity observed within PD-1^+^, CD39^+^ and CD103^+^ TIL populations.

### Both CD4^+^ and CD8^+^ TILs express markers of antitumor reactivity

Having analyzed co-expression and effector profiles of tumor-specific marker expressing populations within overall CD3^+^CD45^+^ TILs, we next examined the biomarker profiles of CD4^+^ or CD8^+^ TIL subsets in ovarian cancer samples. There were more CD4^+^ TILs (mean = 49.6%, 95% CI=43.43 to 55.86) than CD8^+^ TILs (p-value = 0.02, mean = 34.9%, 95% CI=27.88 to 41.85) (**Figure 5A**). CD137 expression was similar in the CD4^+^ and CD8^+^ TIL subsets, suggesting that both CD4^+^ and CD8^+^ T cells are important to antitumor activity in ovarian cancer. More CD8^+^ TILs expressed CD103 (*p*-value <0.001) and CD69 (*p*-value = 0.01). A higher percentage of CD4^+^ TILs expressed CD25 (*p*-value <0.001) and OX40 (*p*-value = 0.01), but there was no significant differences in CD39 or PD-1 expression (**Figure 5B**). When comparing effector molecule expression, an increased frequency of IFNγ (*p*-value = 0.04), TNFα (*p*-value = 0.01), and IL-2 (*p*-value = 0.01) expressing cells was detected in CD4^+^ TILs. CD8^+^ TILs contained greater frequencies of cells expressing Granzyme B (p-value = 0.03), and no difference was detected in perforin expression (**Figure 5C**). These results support the notion that both CD4 and CD8^+^ TILs can express effector molecules, which can be divergent and together may play integral roles in immune responses against tumor cells.

**Figure 5.**
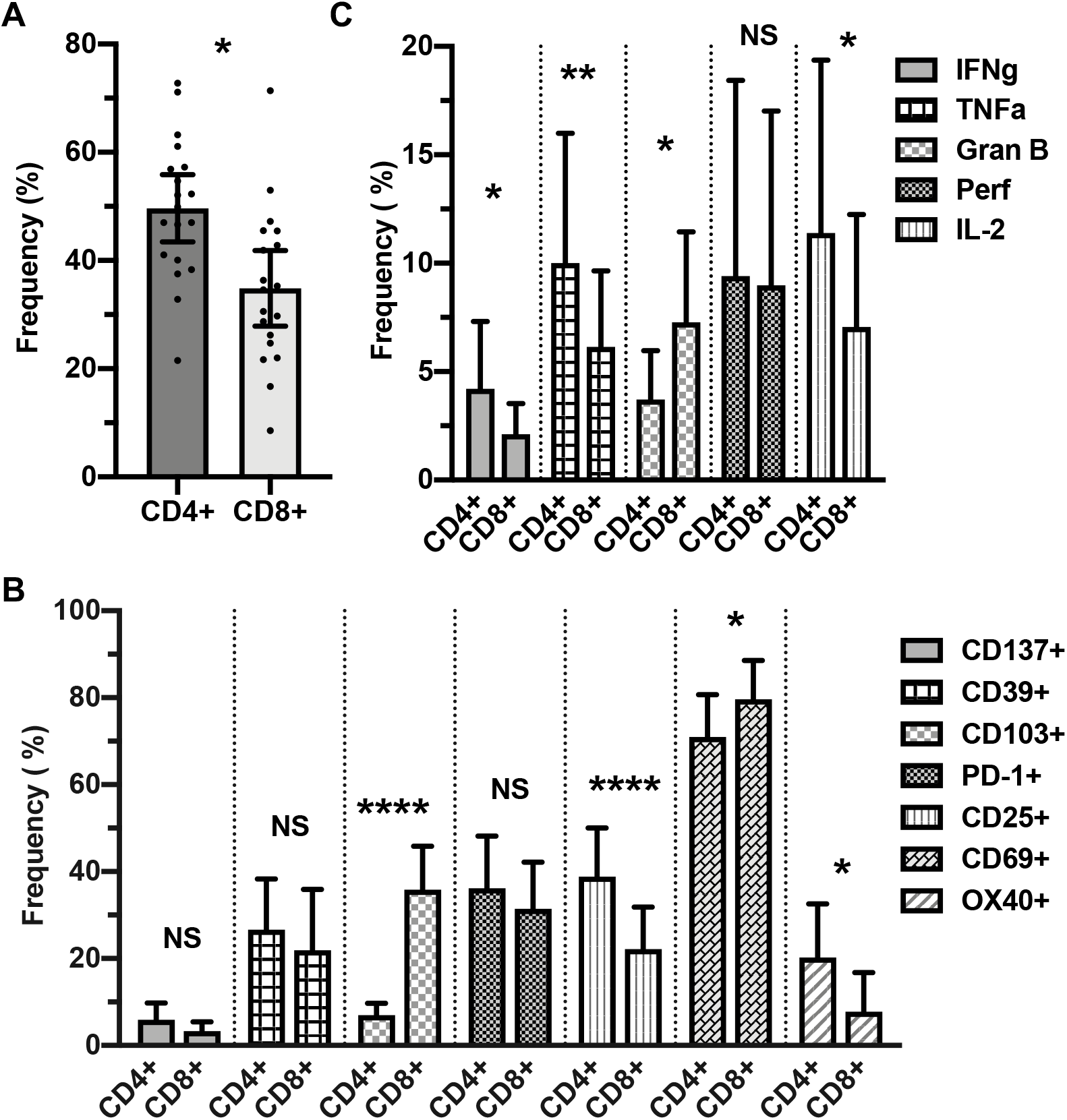

### Tumor-specific TILs display a phenotype indicative of restorable exhaustion

Since our findings indicate that CD137^+^ TILs express effector molecules and other molecules indicative of activation, we queried whether these tumor-specific TILs displayed features of exhaustion, which is commonly associated with chronic tumor-antigen stimulation^20^. We examined to what degree TILs in meta-cluster 5 (MC5) (**Figure 1**,**2**), which harbored the highest frequency of CD137^+^ and effector molecule expressing TILs, exhibit hallmarks of exhaustion. The MC5 population expressed multiple markers indicative of activation and/or exhaustion (**Figure 6A**). MC5 TILs expressed PD-1, albeit at overall lower levels than MC6. MC5 TILs uniquely co-expressed PD-1 and the costimulatory molecule CD28, whose signaling is required for rescue of CD8^+^ T cell activity in anti-PD-1 therapy for cancer ^21^. Activation/exhaustion associated marker expression in CD137^+^ TILs was compared to other TIL populations by examining TIGIT, EOMES, and CD39 expression in CD137^+^ or CD137^-^subsets. CD137^+^ TILs expressed higher levels of the exhaustion-associated markers TIGIT (*p*-value <0.001), EOMES (*p*-value <0.001), and CD39 (*p*-value <0.001), compared to CD137^-^TILs (**Figure 6B**). Since the aforementioned markers can be upregulated by both activated and exhausted T cells, we assessed whether CD137^+^ TILs are skewed towards a EOMES^hi^T-bet^dim^ phenotype associated with dampened effector functions ^22^ or toward a more functional EOMES^dim^T-bet^hi^ phenotype. CD137^+^ TILs were more skewed towards an EOMES^hi^T-bet^dim^ (*p*-value = 0.004) phenotype than their CD137^-^counterparts, supporting the notion that CD137^+^ TILs are exhausted (**Figure 6C, D**). As CD137^+^ TILs appear exhausted but also harbor tumor-specific TILs that express effector molecules and co-express CD28, our results suggest that CD137^+^ TILs have the greatest potential for reinvigoration ^21^.

**Figure 6.**
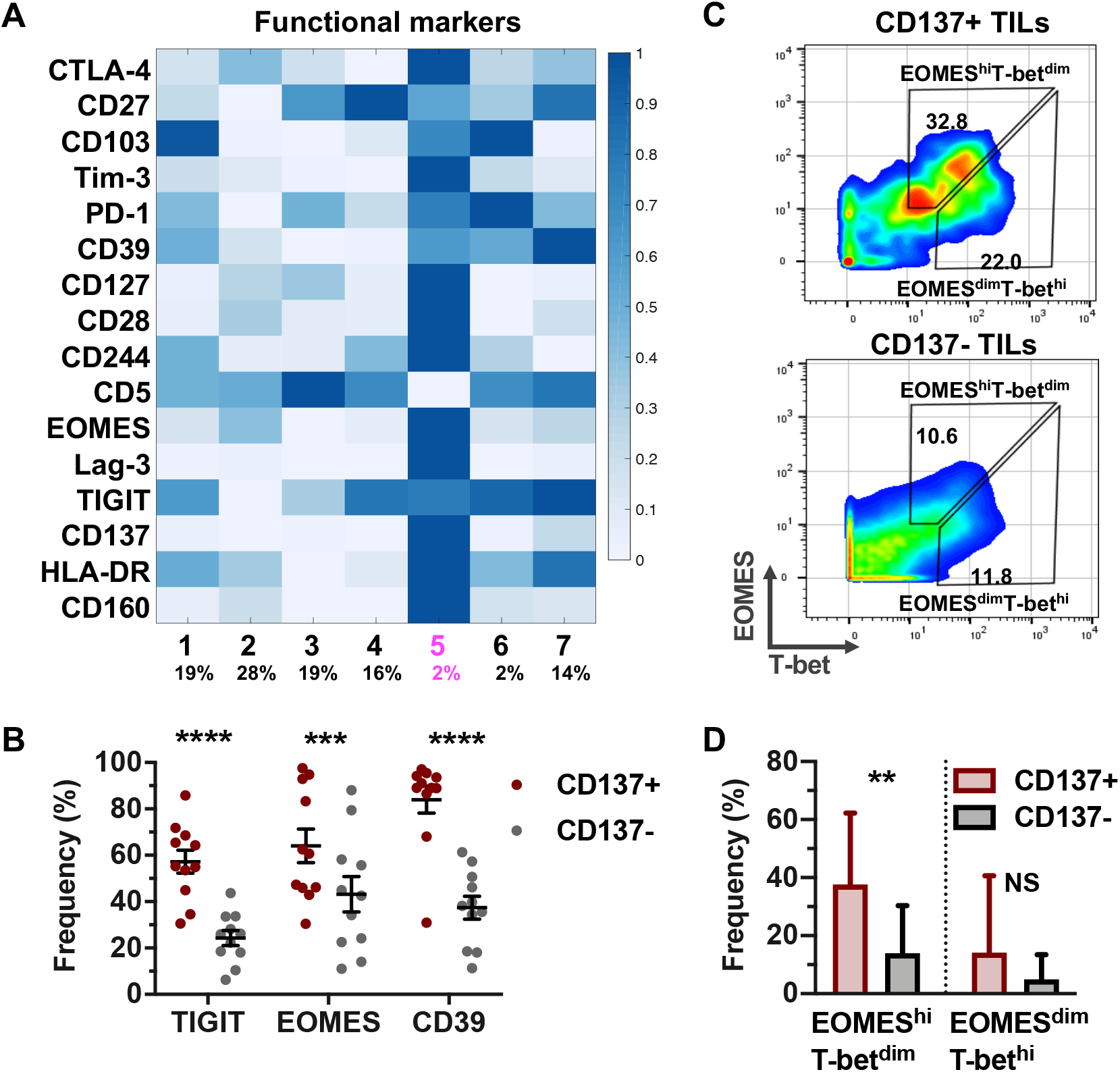

## Discussion

TILs are a heterogeneous population of immune cells that can differ in specificity, differentiation, and function. Biomarkers that identify endogenous tumor-specific TIL subsets are fundamental to immunobiology research, studying mechanisms of endogenous antitumor immunity, isolating tumor-specific T-cell receptors, and optimizing cellular therapies ^5–8^. We observed that TILs expressing effector molecules often co-expressed other biomarkers used to identify tumor-specific TILs. Earlier studies of TILs expressing a single biomarker reported levels of secondary biomarker co-expression, but a direct comparison between various biomarker-expressing TIL subsets had yet to be conducted ^10–13^. We found that a small subset of PD-1^+^, CD103^+^, and CD39^+^ TILs reproducibly co-express CD137. In contrast, most CD137^+^ TILs highly co-express the aforementioned biomarkers, and preferentially express effector molecules, indicating that CD137 more selectively identifies tumor-specific TILs. Further, removing CD137^+^ TILs from other biomarker-expressing TIL subsets reduced their functional activity in response to autologous tumor stimulation, indicating that while PD-1, CD103, and CD39 markers can be used to identify tumor-specific TILs, CD137 expression is a more discriminatory tumor-specific TIL biomarker.

The finding that CD137 expression is a highly selective marker for endogenous tumor-specific TIL identification is supported by previous findings from our lab^10^ [9], and later studies that used CD137 to enrich tumor-specific TILs ^7,10,23^. Our findings contradict results reported by Gros and colleagues showing that both PD-1^+^ and CD137^+^ TIL subsets were tumor-reactive but with PD-1 better identifying tumor-reactive T cells ^11^. Interesting, activation induced expression of CD137 was used to define tumor-reactivity in many of the assays used. The discrepancy between our findings and those reported by Gros et al. may be explained by differences in the cancer type studied as well as the methodology applied. Gros et al. solely focused on CD8^+^ TILs and did not include CD4^+^ TILs. In contrast, the present study, and our previous study that first defined CD137 as a biomarker for tumor-specific TILs ^10^ included CD4^+^ TILs in the analysis. This alone does not account for the discrepancy, since CD137 still served as a better biomarker for tumor-specific CD8^+^ TILs. Identifying endogenous tumor-antigen specific TIL biomarkers in patients has been heavily CD8^+^ T-cell-centric ^11–13,24^, but there is growing appreciation for the role of CD4^+^ T cells in promoting antitumor immunity and immunotherapy efficacy ^25–28^. This is emphasized by our findings that CD4^+^ TILs dominate the ovarian tumor digest environment and have equivalent expression of CD137 as CD8^+^ TILs. Also, with the exception of Granzyme B, CD4^+^ TILs had either equivalent or greater positivity for IFNγ, TNFα, perforin, and IL-2. Our results support the idea that both CD8^+^ and CD4^+^ TILs have integral roles in driving antitumor immune responses and may have divergent antigen-specific responses.

A separate study by Duhen et al. demonstrated that co-expression of CD39 and CD103 TILs can identify tumor-specific TILs within solid tumors. Similar to Gros, *et al*., the work focused on CD8^+^ TILs^13^. Supporting our finding that CD137^+^ TILs often co-express other commonly used tumor-specific TIL biomarkers, both Gros *et. al* and Duhen *et. al*, used CD137 upregulation as a measure to assess tumor-cell recognition by PD-1^+^ or CD39^+^CD103^+^ CD8^+^ TILs in co-culture experiments. Notably, only a subset of enriched PD-1^+^ or CD39^+^CD103^+^ CD8^+^ TILs upregulated CD137 expression after autologous tumor recognition. Unlike Gros *et. al* and Duhen *et. al* studies, we examined TILs from tumor digests without addition of cytokines, establishment of T cell clones, or bulk-expansion. It bears consideration that this methodology can better preserve TIL natural reactivities to autologous tumor antigens with minimal manipulation of TIL biomarker expression.

Immune checkpoint blockade has shown great promise in numerous solid tumors, and successful antitumor responses are thought to rely upon reinvigorated responses by tumor-specific T cells ^20,29,30^. The phenotypic profile of CD137^+^ TILs suggests that they have potential for reinvigoration via checkpoint blockade. CD137^+^ TILs highly expressed multiple co-inhibitory receptors, including PD-1, and were skewed towards a phenotype characteristic of exhausted T cells ^22^ and co-expressed CD28. Expression of CD28 by CD137^+^ TILs is important because restoring exhausted T cell function is dependent on CD28 co-stimulation ^21,31^. However, many cancers, including ovarian cancer, have low response rates to PD-1/PDL1 blockade ^32^. Our data may suggest that one potential explanation is that most patients have too few CD137^+^ TILs to reinvigorate for an effective antitumor response. It is intriguing to hypothesize that the response rate to PD-1/PDL1 blockade may be increased by promoting CD28 signaling to TILs, such as through CTLA-4 blockade. Both CTLA-4 and CD28 bind to CD80 and CD86 on antigen-presenting cells, but CTLA-4 binds CD80 and CD86 with greater affinity and avidity than CD28, enabling it to outcompete CD28 for these ligands. The response rate to anti-PD-1 antibody treatment in ovarian cancer nearly triples when a CTLA-4 blocking antibody is added to the treatment regimen ^37^. Furthermore, agonizing CD137 may aid in promoting antitumor responses in patients, and although CD137 agonism in the clinic has had toxicities ^33,34^, dual bispecific antibodies that agonize CD137 are being developed in order to enhance T cell proliferation and antitumor activity in human cancer without the safety limitations observed in the clinic ^35,36^. The recently developed CD137/OX40 bispecific antibody^35^ may be promising to test in an ovarian cancer model, as we observed that effector molecule expressing CD137^+^ TILs also co-expressed OX40 (**Figure 2A**). Future studies are needed to determine if CD137^+^ TILs are reinvigorated by anti-PD-1 therapy, whether they require CD28 signaling, and how they contribute to successful immune checkpoint blockade monotherapy or combinatorial immunotherapy strategies^35,36^.

Collectively, this work clarifies the differential expression of biomarkers for tumor-specific TILs and demonstrates that CD137 is a more selective biomarker for identifying naturally occurring, tumor-specific TILs than PD-1, CD103, or CD39 within human tumors. We acknowledge there are limitations to this analysis. Our study entirely used ovarian cancer specimens, and results may differ in other cancer types. Also, due to limited cell numbers, we were unable to independently test CD4^+^ and CD8^+^ TILs for TIL subset reactivity, or test restorable exhaustion on PD-1^+^ T cells. Furthermore, PD-1 blockade has low efficacy in *in vitro* assays, and would require sophisticated *in vivo* models and large cell numbers. Nevertheless, we conclude that this work disentangles the differential expression of tumor-specific biomarkers by TILs and identifies CD137 is an ideal singular biomarker for identifying tumor-specific TILs, which provides a deeper understanding of human TILs that may pave a route towards improving immunotherapeutic strategies for cancer.

## Authors’ Contributions

Conceptualization, methodology, project administration, writing, M.A.E. and D.J.P; Supervision and resources, D.J.P. Funding Acquisition. D.J.P and J.C. Formal analysis, visualization, investigation, validation, M.A.E.; Resources and writing, D.K.O.; Writing-review & editing, J.C.

## Acknowledgements

The authors acknowledge the Ovarian Cancer Translational Center for Excellence in the Abramson Cancer Center for support of tumor banking operations through the Tumor BioTrust Collection. The authors thank Dr. Bertram Bengsch and the E. John Wherry laboratory at the University of Pennsylvania, for aid in the development of CyTOF antibody panels and insights on data analysis, as well Takuya Ohtani of the Upenn CyTOF Core for running mass cytometry samples. The authors also acknowledge the personnel of the Wistar Institute’s Flow Cytometry Facility and the Flow Cytometry Core Laboratory at the Children’s Hospital of Philadelphia Research Institute for their aid and expertise in sorting cells.

## Abbreviations

(TILs): tumor-infiltrating lymphocytes
(CI): confidence interval

