## Supplementary figures and images for "Systematic analysis of CD39, CD103, CD137 and PD-1 as biomarkers for naturally occurring tumor antigen-specific TILs"

### Supplement_MAE_20210329 copy.pdf

#### Supplementary Figure 1

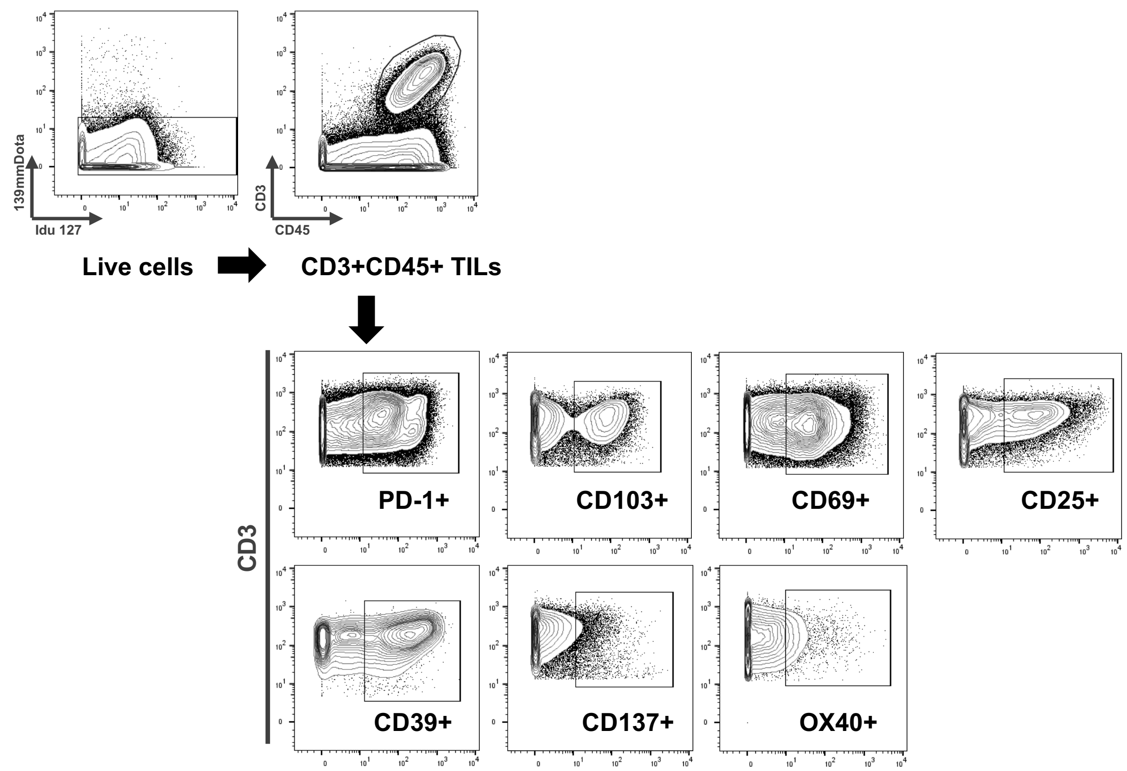

#### Supplementary Figure 2

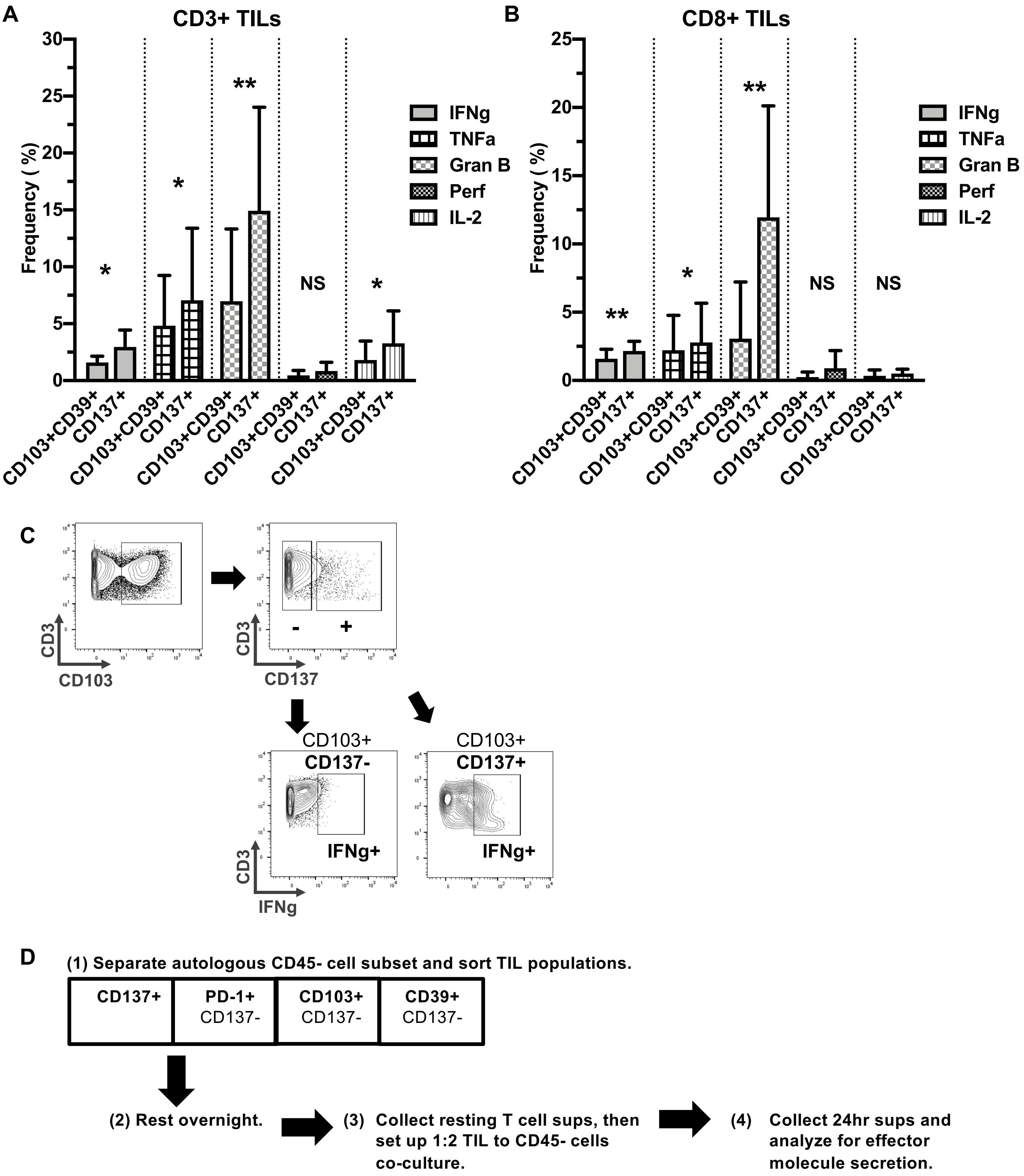
